# An experimental workflow identifies nitrogenase proteins ready for expression in plant mitochondria

**DOI:** 10.1101/2019.12.23.887703

**Authors:** S. Okada, C. M. Gregg, R. S. Allen, A. Menon, D. Hussain, V. Gillespie, E. Johnston, K. Byrne, M. Colgrave, C. C. Wood

## Abstract

Industrial nitrogen fertilizer is intrinsic to modern agriculture yet expensive and environmentally harmful. We aim to reconstitute bacterial nitrogenase function within plant mitochondria to reduce nitrogen fertilizer usage. Many nitrogen fixation (Nif) proteins are required for biosynthesis and function of the mature nitrogenase enzyme, and these will need to be correctly processed and soluble within mitochondria as a pre-requisite for function. Here we present our workflow that assessed processing, solubility and relative abundance of 16 *Klebsiella oxytoca* Nif proteins targeted to the plant mitochondrial matrix using an Arabidopsis mitochondrial targeting peptide (MTP). The functional consequence of the N-terminal modifications required for mitochondrial targeting of Nif proteins was tested using bacterial nitrogenase assays. We found that despite the use of the same constitutive promoter and MTP, MTP::Nif processing and relative abundance in plant leaf varied considerably. Assessment of solubility for all MTP::Nif proteins found NifF, M, N, S, U, W, X, Y and Z were soluble, while NifB, E, H, J, K, Q and V were mostly insoluble. Although most Nif proteins tolerated the N-terminal extension as a consequence of mitochondrial processing, this extension in NifM reduced nitrogenase activity to 10% of controls. Using proteomics, we detected a ∼50-fold increase in the abundance of NifM when it contained the N-terminal MTP extension, which may account for this reduction seen in nitrogenase activity. Based on plant mitochondrial processing and solubility, and retention of function in a bacterial assay, our workflow has identified that NifF, N, S, U, W, Y and Z satisfied all these criteria. Future work can now focus on improving these parameters for the remaining Nif components to assemble a complete set of plant-ready Nif proteins for reconstituting nitrogen fixation in plant mitochondria.

## 1 Introduction

Industrial nitrogen fixation has had a major contribution towards the Green Revolution, and subsequent unprecedented population growth (Smil, 1999). However, the increase in global use of synthetic nitrogen fertilizer has resulted in environmental pollution, contributing to algal blooms, greenhouse gas accumulation and the acidification of soil and water sources (Glibert et al., 2014; Vitousek et al., 1997). There have been several efforts in the past 50 years to look for alternative, more sustainable means to deliver reduced nitrogen to crops, including the use of artificial symbiosis and commensal free-living bacteria (Santi et al., 2014; Oldroyd and Dixon, 2014, Curatti and Rubio, 2014). More recently, advances in synthetic biology have reignited the possibility of generating transgenic crops that can fix their own nitrogen via direct engineering of nitrogenase (Nif) proteins into plants.

Nitrogenase is the enzyme that catalyses biological nitrogen fixation, i.e. the conversion of atmospheric nitrogen to ammonia, and is found exclusively in bacteria and archaea. The molybdenum-dependent nitrogenase consists of two proteins, which are highly oxygen-sensitive: The MoFe protein, a heterotetramer of NifD and NifK, and the Fe protein, a homodimer of NifH. NifDK is the catalytic centre and contains the iron-molybdenum cofactor ([MoFe_7_S_9_C]:homocitrate, FeMo-co; Einsle et al., 2002; Spatzal et al., 2011) and the P-cluster ([Fe_8_S_7_]; Kim and Rees, 1992). NifH is the obligate electron donor, and contains a [Fe_4_S_4_]-cluster (Jang et al., 2000). In addition to the structural proteins, several other Nif proteins are involved in the maturation of the enzyme and assembly of the metalloclusters. These include, but may not be limited to NifB, E, M, N, Q, S, U, V, W, X, Y, and Z (Ribbe et al., 2014). Furthermore, nitrogenase also utilizes specific electron transport proteins, NifF and NifJ (Deistung et al., 1985; Shah et al., 1983). For optimum nitrogenase activity, the stoichiometry of the numerous Nif proteins and their temporal expression needs to be tightly regulated (Pozza-Carrion et al., 2015).

The mitochondrial matrix has been shown to be a suitable location to express some of the most oxygen-sensitive Nif proteins in a functional form (Lopez-Torrejon et al., 2016; Buren et al., 2017a, Buren et al., 2019). However, it is currently technically difficult to directly introduce transgenes into the mitochondrial genome and recover stable transgenic plants (Macmillan et al., 2019). In this study we rely upon the endogenous mitochondrial protein transport pathway for the expression of nuclear-encoded genes within the mitochondrial matrix (Fig. 1; reviewed by Murcha et al., 2014). This process involves the use of mitochondrial targeting peptides (MTPs) as translational fusions at the N-terminus of each Nif protein (MTP::Nif). After translation in the cytoplasm the MTP::Nif protein is actively transported to the mitochondrial matrix through the outer and inner transmembrane import complexes. The MTP is cleaved within the mitochondrial matrix by the mitochondrial processing peptidase (MPP) and the remaining C-terminus is folded into the mature protein. The MPP-dependent processing of the MTP results in residual amino acids at the N-terminus of the transgenic Nif proteins, and here we term this the ‘scar’ peptide.

**Fig. 1:**
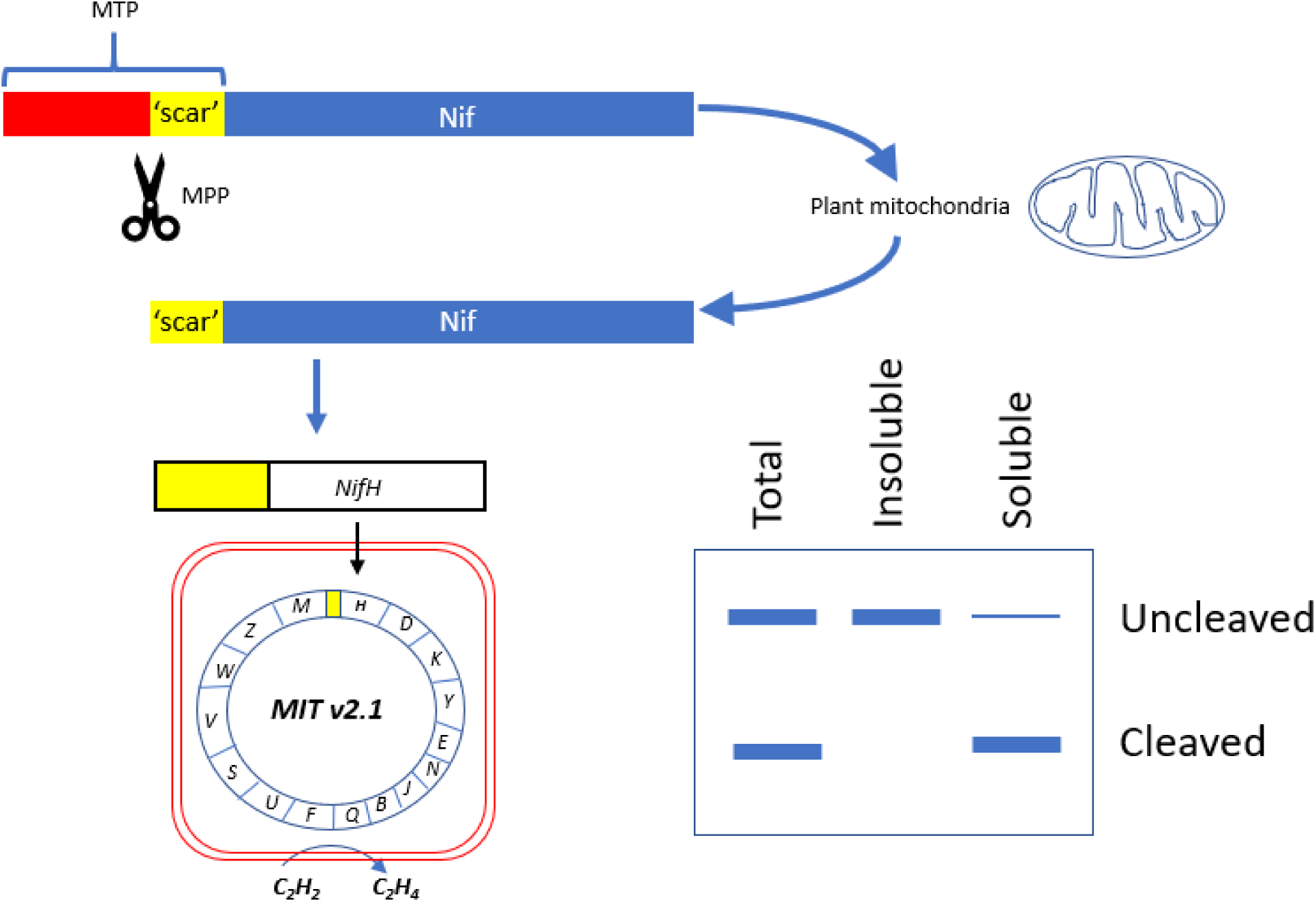
Schematic of the workflow to assess key features of Nif proteins targeted to plant mitochondria. Translational fusions of the MTP to Nif proteins are expressed in *N. benthamiana* leaf to test preprotein processing and solubility. Scar::Nif protein fusions are expressed in *E. coli* for function testing by acetylene reduction. MTP, ‘scar’ peptide, Nif not to scale. MTP, mitochondrial targeting peptide; MPP, mitochondrial processing peptidase.

These N-terminal modifications could potentially impact the function of Nif proteins, and it is currently unknown if all Nif proteins can tolerate a scar peptide. The clearest example of scar peptides being tolerated was shown by the isolation of functional NifH from yeast mitochondria, a result dependent on the import, MPP processing and refolding of MTP::NifH and MTP::NifM (Lopez-Torrejon et al., 2016). As an alternative approach to using eukaryotic platforms that currently present numerous challenges, bacterial-based assays can be used to guide modifications to Nif proteins (Yang et al., 2018).

Another important consideration for function is the solubility of each Nif protein in plant mitochondria. Burén et al. (2017a) found that NifB from *Azotobacter vinelandii* was insoluble when it was targeted to the mitochondrial matrix of both yeast and plants. Encouragingly these authors found two variants of NifB from a thermophilic archaea that was soluble within the yeast mitochondrial matrix and active in a reconstitution assay (Buren et al., 2017a, 2019). Aside from NifB, the solubility of other Nif proteins has not been directly assessed within plant mitochondria.

Previously we targeted 16 *Klebsiella oxytoca* Nif proteins to plant mitochondria using an MTP of 77 amino acids. In this study, we design and test a shorter MTP fused to the 16 Nif proteins and assess abundance, processing and solubility of the synthetic proteins when targeted to the plant mitochondria. This shortened MTP is cleaved within the matrix to leave a nine amino acid scar, and we use a bacterial assay to assess the functional impact of this N-terminal modification to each Nif protein. Our analysis has identified a subset of MTP::Nif proteins that satisfy the requirements of being soluble, correctly processed, and functional. These MTP::Nif proteins are now ready for further downstream analysis and functional testing in plant mitochondria.

## 2 Results

### 2.1 Design and testing a 51 amino acid MTP for MPP processing of MTP::Nif proteins in plant mitochondria

Our previous work targeting Nif proteins to the mitochondrial matrix utilised a 77 amino acid (AA) peptide of the N-terminus of the ATP synthase γ subunit from *Arabidopsis thaliana* (pFAγ77, Allen et al., 2017). Processing of pFAγ77 by the MPP resulted in a 35 AA residual ‘scar’ on the N-terminus of the Nif protein. However, introducing long N-terminal extensions to Nif proteins may impair function via steric hindrance. We therefore wanted to shorten the MTP sequence to minimise the remaining scar yet retain targeting capability to the plant mitochondrial matrix. Previously Lee et al. (2012) showed that residues 52 to 77 of the original pFAγ77 were possibly not required for transporting and processing of green fluorescent protein to the mitochondrial matrix. Based on these observations we redesigned a shorter MTP with a length of 51 AA, here termed pFAγ51. pFAγ51 is predicted to leave a nine AA N-terminal extension after MPP processing that we term scar9 (amino acid sequence ISTQVVRNR; Fig. 2A; Huang et al., 2009). To confirm the site of cleavage of pFAγ51 fused to Nif proteins we constructed a translational fusion of pFAγ51::NifU::Twin-Strep-tag® (SN166; Twin-Strep-tag®, Schmidt et al., 2013) using a modular Golden Gate assembly method (Weber et al., 2011). After infiltration of SN166 into *N. benthamiana* leaf this protein was purified by affinity chromatography (Suppl. Fig. 1) and subjected to proteomics analysis, which identified ISTQVVR as the N-terminal peptide sequence. This result confirmed that the shortened MTP, pFAγ51, was functional and processed as predicted to leave a nine AA scar at the N-terminus.

**Fig. 2:**
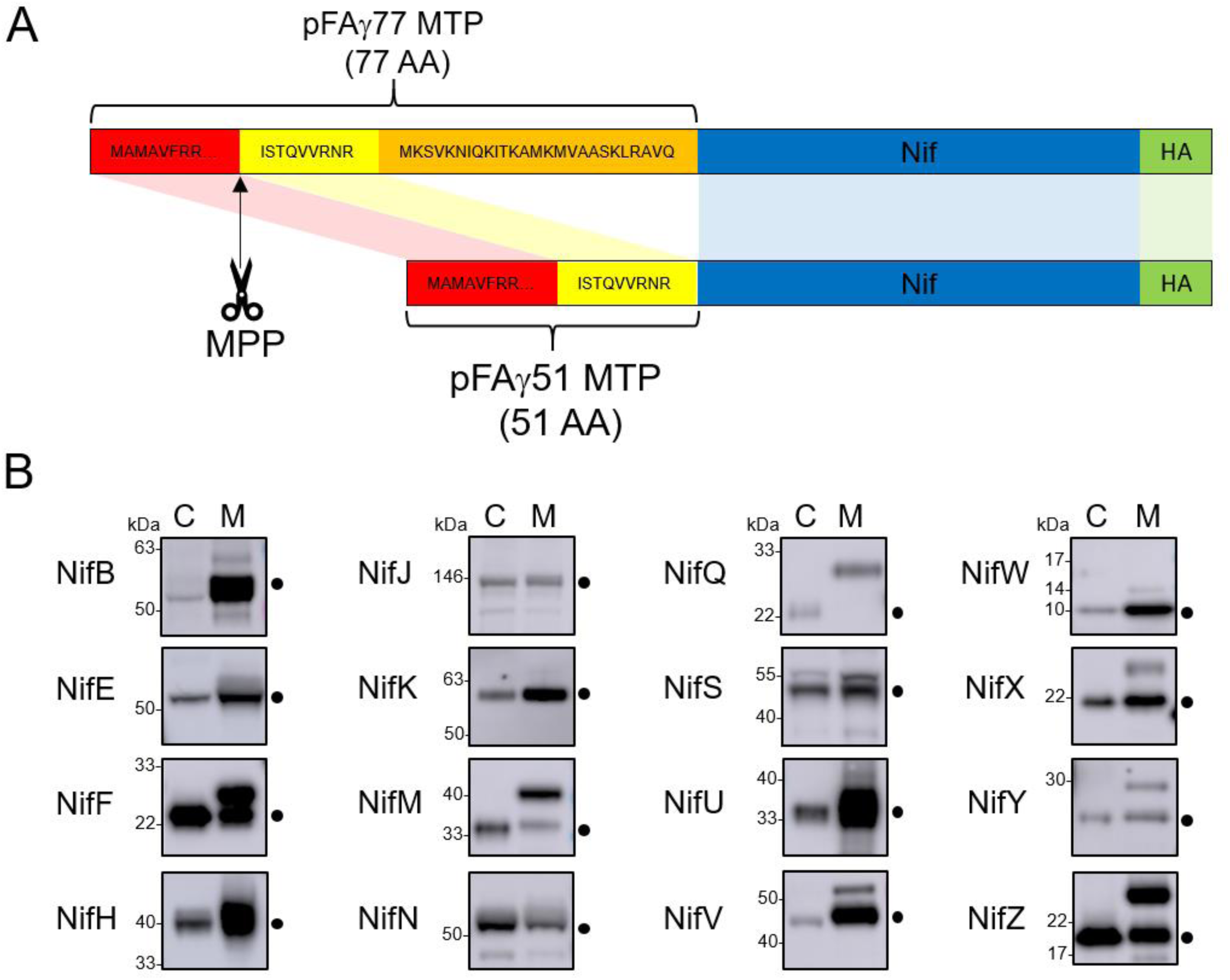
Design of the pFAγ51 MTP and assessment of its cleavage when fused to different Nif proteins expressed in plants. **(A)** Schematic of the truncation of the N-terminal amino acid sequence of the *A. thaliana* F1 ATPase γ subunit from 77 to 51 residues. The 26 amino acid residues in the orange section of pFAγ were removed. MTP, Nif, HA epitope tag are not to scale. MTP, mitochondrial targeting peptide; MPP, mitochondrial processing peptidase. **(B)** Western blot analysis (α-HA) of individual pFAγ51::Nif::HA, pFAγ51::HA::NifK, 6×His::Nif::HA and 6×His::HA::NifK proteins transiently expressed in *N. benthamiana* leaf. Black dots point to the size of the correctly processed pFAγ51::Nif::HA protein. C, cytoplasmic expression; M, mitochondrial targeted. Panels of individual Nif proteins shown in B were extracted from full blot images presented in Suppl. Fig. 2.

### 2.2 Most pFAγ51::Nif proteins are targeted to and processed in the plant mitochondrial matrix

We next wanted to assess whether the shortened MTP, pFAγ51, was able to target other Nif proteins to the plant mitochondrial matrix. We generated expression constructs for 16 Nif proteins with translational fusions of pFAγ51 at the N-terminus and a HA epitope tag at the C-terminus, resulting in the generic structure pFAγ51::Nif::HA (Fig. 2A, plant expression constructs listed in Suppl. Table 2). NifK was the only protein for which a different construct was made, where the HA-tag was included at the N-terminus, since any C-terminal fusions render NifK non-functional (Yang et al., 2018). Expression of pFAγ51::NifD::HA will be reported as part of a separate study (Allen et al., 2019 https://doi.org/10.1101/755116) and therefore was not included in this study. We constructed control expression plasmids to mimic the processed protein size for all Nif proteins by replacing pFAγ51 with 6×His. These 6×His::Nif::HA proteins are expected to be located to the cytoplasm. Both pFAγ51::Nif::HA and 6×His::Nif::HA were infiltrated, separately, in transient *N. benthamiana* leaf assays and the migration speeds of the expressed proteins assessed by Western blot analysis.

Comparison of each Nif protein targeted to either the mitochondrial matrix or cytoplasm demonstrated that 15 of 16 Nif proteins targeted to the mitochondria using pFAγ51 were processed by MPP, although with variable efficiency (Fig. 2B, full blot images provided in Suppl. Fig. 1). We observed three general classes of processing efficiency – the first being efficiently cleaved, second being partially cleaved, and the last being poorly cleaved to not cleaved at all. Efficient cleavage was found for eight pFAγ51::Nif::HA (NifB, E, H, J, N, U, V, W) and pFAγ51::HA::NifK proteins. Six pFAγ51::Nif::HA proteins (NifF, M, S, X, Y, Z) were partially cleaved, as evidenced by the presence of two HA-dependent signals, the faster band migrating at the speed of the corresponding 6×His::Nif::HA control and another slower band running at a speed consistent with the size of the unprocessed pFAγ51::Nif::HA (predicted unprocessed protein sizes provided in Suppl. Table 1). The only protein displaying no processing was pFAγ51::NifQ::HA, with only a signal found for a protein consistent with the unprocessed size (Fig. 2B). For some pFAγ51::Nif::HA proteins, e.g. NifB, E, H, S, U and Z there were additional signals at a higher molecular weight, which could arise from dimerization or oligomerization (Suppl. Fig. 2, 3 and 4), similar to what has been previously observed (Allen et al., 2017). In some instances, e.g. NifJ, we also observed degradation products (Suppl. Fig. 2).

Using Western blot analysis to assess MPP processing further provided an indication of the relative abundance of each Nif protein in the transient leaf assay system. In general, we found that most of the pFAγ51::Nif::HA and pFAγ51::HA::NifK proteins were readily detectable, although their abundance varied (Fig. 2B). For instance pFAγ51::NifY::HA had a relatively low signal intensity whereas NifH, F and U had the highest signal intensities. Another observation worth noting was the difference in abundance between cytoplasmic- and mitochondria-targeted Nif proteins. NifB, E, H, U, V and W proteins targeted to the mitochondria accumulated to higher levels than when targeted to the cytoplasm, whereas the level of expression of the other Nif proteins were approximately equal between the mitochondrial and cytoplasmic forms.

### 2.3 Solubility testing of Nif proteins targeted to leaf mitochondria

As all Nif proteins need to be soluble for function we tested solubility of the Nif proteins when targeted to the plant mitochondrial matrix. Total protein extracts of *N. benthamiana* leaf tissue individually expressing 16 pFAγ51::Nif::HA and pFAγ51::HA::NifK were separated into soluble and insoluble fractions and analyzed by Western blot (Fig. 3, full blot image provided in Suppl. Fig. 3). We found that the relative abundance of correctly processed Nif proteins in the soluble fraction varied for each pFAγ51::Nif::HA/pFAγ51::HA::NifK. For example NifF, M, N, Q, S, and W were predominantly in the soluble fraction in both the correctly processed and unprocessed form. Conversely NifB, E, H, J, and V were not found in the soluble fraction, despite being correctly processed. As NifM may be required for stability and solubility of NifH in bacteria (Lei et al., 1999; Howard et al., 1986) we tested coexpression of mitochondrially targeted pFAγ51::NifH::HA and pFAγ51::NifM::HA in transient leaf assays but found no improvement in the solubility of NifH (data not shown). We found a third band for NifF between the processed and unprocessed form, which was possibly an artefact of protein extraction, as it was not detected in Western blot of total protein. Interestingly, pFAγ51::NifQ::HA produced a faint band approximately the size of the correctly processed form in the soluble fraction, which was not detectable in the total protein Western blot. To assess if atmospheric oxygen affected Nif protein solubility the same 16 pFAγ51::Nif::HA/pFAγ51::HA::NifK proteins were isolated from infiltrated plants under anaerobic conditions and subjected to Western blot analysis. We found that anaerobic conditions during protein extraction did not change their solubility (Suppl. Fig. 4).

**Fig. 3:**
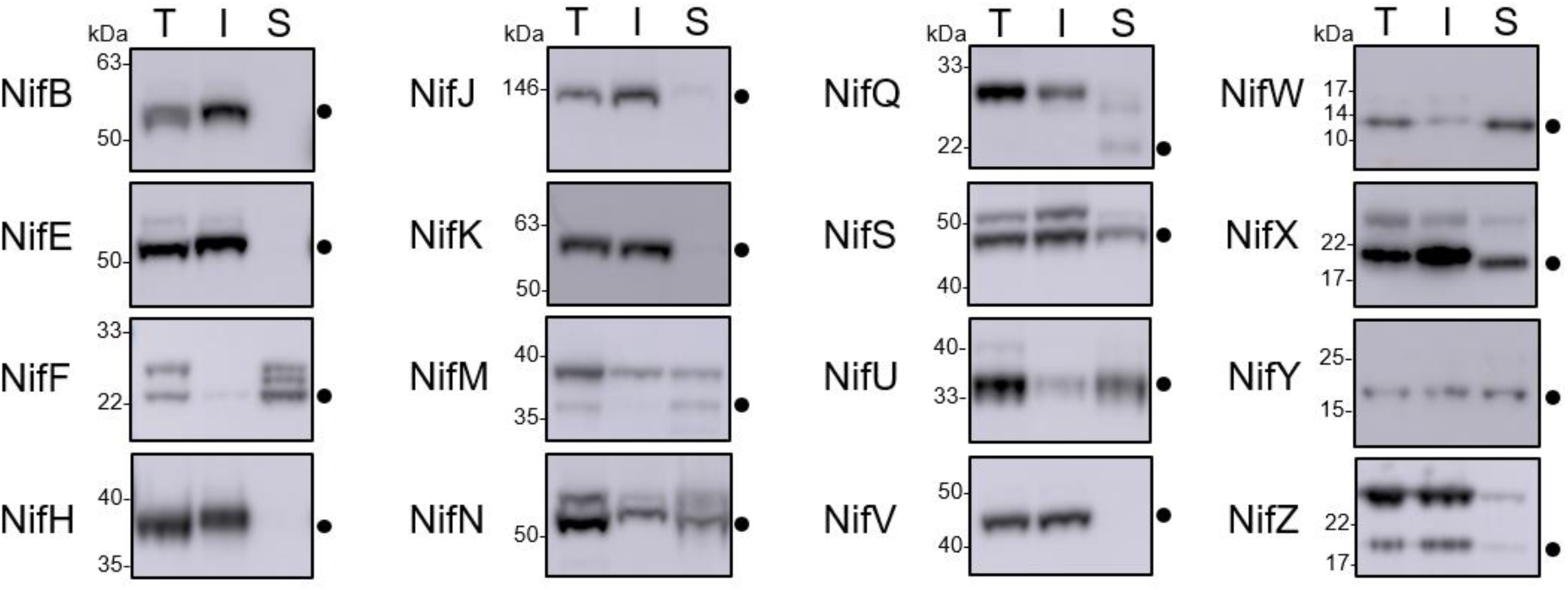
Assessment of the solubility of pFAγ51::Nif::HA and pFAγ51::HA::NifK proteins in plants. Western blot analysis (α-HA) of individual pFAγ51::Nif::HA and pFAγ51::HA::NifK proteins transiently expressed in *N. benthamiana* leaf. Black dots point to the size of the processed pFAγ51::Nif::HA and pFAγ51::HA::NifK protein. T, total protein; I, insoluble fraction; S, soluble fraction. Panels of individual Nif proteins shown were extracted from full blot images presented in Suppl. Fig. 3.

### 2.4 Testing function of modified Nif proteins with an N-terminal extension

Using a bacterial assay we tested the functional impact of adding nine AA to the N-terminus of each Nif protein, which mimics the residual scar peptide that remains after MPP processing of pFAγ51::Nif in plant mitochondria. We adopted the MIT v2.1 plasmid system (Smanski et al., 2014) for this assay, and fused the nine AA scar9 sequence, MSTQVVRNR, to the N-terminus of each Nif protein, mimicking the length and sequence of pFAγ51 after MPP cleavage. An example of the process is outlined with scar9::NifH replacing NifH within MIT v2.1 (Fig. 4). Each scar9::Nif was tested individually in separate MIT v2.1 plasmids in the same manner (bacterial expression constructs listed in Suppl. Table 3). It is worth noting that MIT v2.1 does not have NifX and therefore we did not test the impact of scar9 on this protein with respect to nitrogenase function. As a negative control we removed NifH, D, K, Y, E, N, and J from MIT v2.1 and made a non-functional plasmid, here termed ‘pB-ori’. As further controls we made other modifications, such as adding a HA epitope tag to the C-terminus of NifK, namely NifK::HA, or removing NifM from MIT v2.1 (cf. Lei et al., 1999; Howard et al., 1986), both of which resulted in the expected loss of nitrogenase function (Table 1). Function testing of the individual scar9::Nif proteins in *E. coli* by acetylene reduction showed that activity was retained for all 16 scar9::Nif proteins although there was variation in activity levels. Notably scar9::NifJ had three times the activity of the positive control, and scar9::NifQ, H, B and F were mildly increased (130-150% activity relative to MIT v2.1). In contrast, scar9::NifM only retained approximately 10% activity of the positive control.

**Table 1.** Effect of pFAγ51 nine amino acid ‘scar’ (scar9) peptide translationally fused to individual Nif proteins on nitrogenase function. Values are presented as % acetylene reduction activity compared to MIT v2.1. pB-ori, negative control containing *nifBQFUSVWZM*; Δ*nifM*, NifM coding sequence removed from MIT v2.1; NifK::HA, HA epitope tag fused to the C-terminus of NifK; S.D., standard deviation (n=2-6).

| Construct | % activity of<br>MIT v2.1 | S.D. |
| --- | --- | --- |
| MIT v2.1 | 100 | 18 |
| scar9::NifJ | 309 | 86 |
| scar9::NifQ | 158 | 22 |
| scar9::NifH | 148 | 15 |
| scar9::NifB | 144 | 34 |
| scar9::NifF | 131 | 8 |
| scar9::NifD | 110 | 9 |
| scar9::NifW | 95 | 8 |
| scar9::NifV | 81 | 11 |
| scar9::NifU | 80 | 4 |
| scar9::NifY | 65 | 4 |
| scar9::NifK | 60 | 10 |
| scar9::NifN | 46 | 9 |
| scar9::NifE | 42 | 1 |
| scar9::NifS | 41 | 1 |
| scar9::NifZ | 37 | 18 |
| scar9::NifM | 9 | 1 |
| $\Delta nifM$ | 6 | 4 |
| NifK::HA | 0 | 1 |
| pB-ori ($\Delta nifHDKENJ$) | 0 | 0 |

**Fig. 4:**
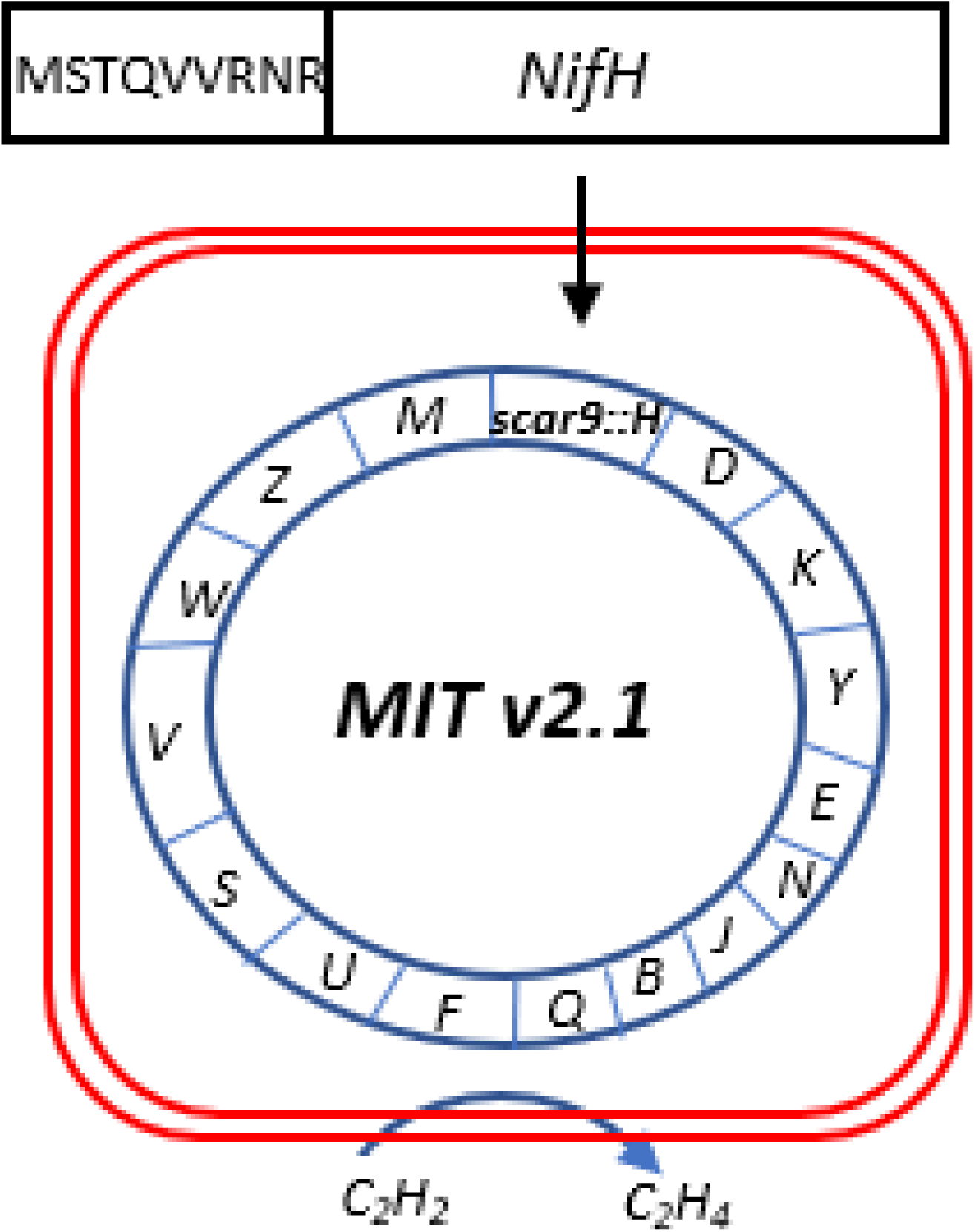
Representation of a modified MIT v2.1 to test function in *E. coli* strain JM109. This example shows a scar9 motif translationally fused to NifH. One scar9::Nif was tested per expression plasmid.

To assess if the scar9 peptide had any influence on the relative abundance of each Nif protein in *E. coli*, we measured the relative protein abundance of the Nif proteins D, K, H, S and M, in *E. coli* containing the native and modified forms of MIT v2.1 using targeted proteomics. We also measured the relative abundance of a peptide specific to chloramphenicol acyltransferase (CAT), the coding nucleotide sequence of which was present in all MIT v2.1 plasmids. We found that signals for NifS and CAT peptides were relatively consistent across all the samples (Suppl. Fig. 5, Suppl. Table 1). As expected, we also found that signals specific to NifM, peptides M-1 and -2, were not found in *E. coli* containing MIT v2.1 in which NifM was deleted, and that signals specific for NifD, K, H, Y, E, and N were not found in *E. coli* with pB-ori, where these genes were removed from MIT v2.1 (data not shown). The most unexpected change was found for NifM, where the relative protein abundance was approximately ∼50-fold higher in *E. coli* expressing scar9::NifM, relative to those expressing other scar9::Nifs (Suppl. Fig. 5).

## 3 Discussion

In this study we present a workflow to assess key functional prerequisites of MTP::Nif proteins targeted to plant mitochondria. These were cleavage of the MTP by the MPP in the mitochondrial matrix, solubility, relative protein abundance and tolerance of N-terminal scar sequences for function. The plant- and bacterial-based assays identified seven Nif proteins, namely NifF, N, S, U, W, Y and Z, that we consider ready for metabolic engineering of nitrogenase into plant mitochondria using pFAγ51 as the MTP. Importantly, we also found other MTP::Nif proteins to be either poorly processed and/or insoluble in plant mitochondria, or impaired in functional assays. The identification of these problematic MTP::Nif proteins can guide targeted improvements in the future.

We have found relative protein abundance, processing efficiency, and solubility of the 16 different MTP::Nif proteins varied, despite use of the same MTP and promoter for each plant expression construct. This variation illustrates how intrinsic properties of each Nif protein influence these attributes in plant mitochondria. Assuming that levels of plant expressed Nif proteins will need to reflect those of diazotrophic bacteria, future studies will need to adjust promoter strength and/or translation rates accordingly. For example NifY was the least abundant in our experiments, and efforts are needed to improve these levels to better mimic those found in naturally occurring systems (Smanski et al., 2014).

Similarly we found some MTP::Nif proteins were poorly cleaved by the MPP, in particular MTP::NifQ. A potential reason for this may be that the preprotein is unable to enter the mitochondrial matrix due to the MTP::Nif protein being resistant to unfolding (Voos et al., 1993, 1996). We also found that most Nif proteins that were successfully cleaved within the mitochondrial matrix tended to accumulate to higher levels relative to their cytoplasmic counterparts, suggesting that mitochondrial processing may stabilize Nif proteins relative to those located in the cytoplasm.

In our experiments we found several MTP::Nif proteins were insoluble in the plant mitochondrial matrix. Notably the key protein NifH from *K. oxytoca* was among this set. Interestingly *A. vinelandii* NifH and NifM when targeted to yeast mitochondria produced a functional Fe protein (Lopez-Torrejon et al., 2016), indicating that both these proteins were sufficiently soluble in yeast. In agreement with prior results, we found that *K. oxytoca* NifB was insoluble when targeted to plant mitochondria, as was described for *A. vinelandii* NifB when targeted to yeast and plant mitochondria (Burén et al., 2017a).

Overcoming these processing, solubility and abundance issues we encountered will require a range of approaches. There is evidence in yeast mitochondria that the import of transgenic cargoes can be improved by the use of longer MTPs (Wilcox et al., 2005). Therefore screening alternate length MTPs may reveal certain MTP::Nif combinations that overcome problems with recalcitrant import. Other bacterial or archaeal variants of the nitrogen fixation pathway can also provide the means to improve targeting, processing and ultimately activity of nitrogenase within mitochondria, as was shown for nifB (Burén et al., 2017a, 2019).

Although this report concentrates on attributes of Nif proteins expressed individually, there may be combinations of Nif proteins that when expressed together improve abundance or solubility. Functional *Azotobacter* Nif proteins have been successfully expressed in combination within yeast mitochondria (Burén et al., 2017b, 2019; Lopez-Torrejon et al., 2016), and these combinations may overcome problems that we report here. Finally, physically linking Nif proteins into larger multi-domain polyproteins (Allen et al., 2019 https://doi.org/10.1101/755116, Yang et al., 2018) that can assist in protein assembly may also overcome problems associated with solubility identified in this report.

The function of nitrogenase may be impacted by residual terminal scar residues remaining after mitochondrial targeting. Yang et al. (2018) demonstrated that the addition of the tobacco etch virus protease cleavage site to the C-terminus of NifK abolished activity, a result that could be predicted from the close interaction of NifK with NifD (Spatzal et al., 2011). Here we tested a 9 AA scar on the N-terminus of 16 Nif proteins and found both positive and negative impacts on overall nitrogenase activity. Importantly we found that the key protein NifH supported nitrogen fixation with the N-terminal addition. NifH has three different functions, firstly, donating electrons to NifDK, secondly, maturation of the [Fe_8_S_7_] P-clusters within NifDK, and thirdly as a molybdenum and homocitrate insertase to NifEN (reviewed in Hu and Ribbe 2013). In a previous study, NifH isolated from yeast mitochondria was capable of donating electrons to NifDK (Lopez-Torrejon et al., 2016). Our study demonstrates that the additional functions of NifH can also occur despite the addition of the MTP scar sequence.

Examples of Nif proteins that did not tolerate the N-terminal extension with the MTP that was tested were NifE, N and M. In the case of NifEN these proteins form a stable heterotetramer, but also interact with numerous other Nif proteins during the biogenesis of FeMo-co, including NifB, NifY, NafY, NifX and NifH (Ribbe et al., 2014). The N-terminal extension on NifE and NifN that was tested in our study may have reduced nitrogenase function via steric hinderance within protein-protein interactions associated with NifEN. The most severe impact on nitrogenase function was found for scar9::NifM (∼10% of control), but in that instance proteomics analysis found that the abundance of scar9::NifM was highly up-regulated compared with other modified MIT v2.1 plasmids. As nitrogenase activity is highly sensitive to changes in Nif protein levels (reviewed in Martinez-Argudo et al., 2004) this misregulation may account for the decrease in nitrogenase activity seen with scar9::NifM, rather than reflecting steric interference.

Although mitochondria are considered potentially suitable to support nitrogenase activity, impediments remain to successfully translocating all Nif proteins to the organelle. This is not surprising considering the large span of evolutionary time separating the emergence of nitrogenase in bacteria from the origins of mitochondria in eukaryotes (Muller et al., 2012; Poole and Gribaldo, 2014). Our testing uncovered some Nif proteins that we consider compatible with translocation to plant mitochondria and other Nif proteins that require further improvement. The experimental framework outlined here can be applied systematically to iteratively improve each Nif protein with the eventual goal of assembling the entire pathway within plant mitochondria.

## 4 Materials and Methods

### 4.1 Construction of plasmids for *Nicotiana benthamiana* leaf transient expression

Plasmids for transient expression in *N. benthamiana* leaf were constructed using a modular cloning system with Golden Gate assembly (Weber et al., 2011). DNA parts as individual plasmids (Thermo Fisher Scientific, ENSA), each containing the 35S CaMV promoter (EC51288), the gene coding for the first 51 amino acids of the Arabidopsis F1-ATPase γ subunit (pFAγ51), plant codon optimised *nifH* (EC38011), *nifK* (EC38015), *nifY* (EC38019), *nifE* (EC38016), *nifN* (EC38024), *nifJ* (EC38022), *nifB* (EC38017), *nifQ* (EC38025), *nifF* (EC38021), *nifU* (EC38026), *nifS* (EC38018), *nifV* (EC38020), *nifW* (EC38027), *nifZ* (EC38029), *nifM* (EC38023), *nifX* (EC38028), plant codon optimised HA epitope tag (EC38003), and CaMV terminator (EC41414) were assembled into plant expression vectors (EC47772, EC47742, EC47751, EC47761, EC47781) using Type IIS restriction cloning. The plasmid ID and descriptions are listed in Supplementary Table 2.

### 4.2 Plant growth and transient transformation of *N. benthamiana*

*N. benthamiana* plants were grown in a Conviron growth chamber at 23°C under a 16:8 h light:dark cycle with 90 μmol/min light intensity provided by cool white fluorescent lamps. *Agrobacterium tumefaciens* strain GV3101 (SN vectors) or AGLI (P19 vector) cells were grown to stationary phase at 28°C in LB broth supplemented with 50 mg/mL carbenicillin or 50 mg/mL kanamycin, according to the selectable marker gene on the vector, and 50 mg/mL rifampicin. Acetosyringone was added to the culture to a final concentration of 100 μM and the culture was then incubated for another 2.5 h at 28°C with shaking. The bacteria were pelleted by centrifugation at 5000 x *g* for 10 min at room temperature. The supernatant was discarded, and the pellet was resuspended in 10 mM MES pH 5.7, 10 mM MgCl_2_ and 100 μM acetosyringone (infiltration buffer) after which the OD_600_ was measured. A volume of each culture, including the culture containing the viral suppressor construct 35S::P19, required to reach a final concentration of OD_600_ = 0.10 was added to a fresh tube. The final volume was made up with the infiltration buffer. Leaves of five-week-old plants were then infiltrated with the culture mixture and the plants were grown for five days after infiltration before leaf samples were harvested for further analysis/experiments.

### 4.3 Western blot analysis of Nif proteins transiently expressed in *N. benthamiana*

To assess the processing of mitochondrially targeted and cytoplasmically located proteins, leaf disks of 180 mm^2^ were harvested from *N. benthamiana* and the proteins were extracted, subjected to SDS-PAGE and Western blot according to Allen et al. (2017). Monoclonal anti-HA antibody produced in mouse (Sigma-Aldrich) was used as the primary antibody and Immun-Star Goat Anti-Mouse (GAM)-HRP conjugate (Bio-Rad) was used as the secondary antibody. The PageRuler™ Prestained Protein Ladder (Thermo Fisher Scientific) and the BenchMark™ Pre-Stained Protein Ladder (Thermo Fisher Scientific), which was re-calibrated against the unstained BenchMark™ protein ladder to 146 kDa, 91 kDa, 63 kDa, 50 kDa, 40 kDa, 33 kDa, 22 kDa, 17 kDa, 14 kDa and 10 kDa, were used as molecular size markers.

For solubility testing the harvested leaf tissue was ground in liquid nitrogen using a mortar and pestle and transferred to a microfuge tube. Three hundred (300) μL of cold solubility buffer (50 mM Tris-HCl pH 8.0, 75 mM NaCl, 100 mM mannitol, 2 mM DTT, 0.5% (w/v) polyvinylpyrrolidone (average MW 40 kDa), 5% (v/v) glycerol, 0.2 mM PMSF, 10 μM leupeptin and 0.5% (v/v) Tween® 20) was added and the samples were centrifuged for 5 min at 16,000 x g at 4°C. The supernatant was transferred to a fresh tube and the pellet was resuspended in 300 μL of fresh cold solubility buffer. Both, the supernatant (sample 1) and the resuspended pellet (sample 2) were centrifuged again for 5 min at 16,000 x *g* at 4°C. From sample 1 a subsample was taken, which is referred to as the soluble fraction. This subsample was mixed with an equivalent amount of 4 x SDS buffer (250 mM Tris-HCl pH 6.8, 8% (w/v) SDS, 40% (v/v) glycerol, 120 mM DTT and 0.004% (w/v) bromophenol blue). After the second centrifugation step, the supernatant of sample 2 was discarded. The pellet is referred to as the insoluble fraction. The pellet was resuspended in 300 μL 4 x SDS buffer and 300 μL of solubility buffer were added. When soluble and insoluble fractions were compared to the amount of total protein, the leaf piece for the total protein sample was ground as described above. However, the ground sample was resuspended in 300 μL 4 x SDS buffer and 300 μL of solubility buffer were added. Samples for the total, insoluble and soluble fractions were heated at 95°C for 3 min and then centrifuged at 12000 x *g* for 2 min. 20 μL of the supernatant containing the extracted polypeptides was loaded on a NuPAGE Bis Tris 4-12% gels (Thermo Fisher Scientific) for gel electrophoresis and Western blot analysis.

For Western blot analysis of anaerobically extracted proteins, the extractions were carried out in an anaerobic chamber (COY Laboratory Products) filled with a H_2_/N_2_ atmosphere (2-3%/97-98%). Anaerobic extraction solutions were prepared at a Schlenk line in a bottle equipped with a butyl rubber septum by at least four cycles of evacuating and purging with N_2_. Leaf disks were ground in cold solubility buffer instead of liquid nitrogen.

### 4.4 Isolation of pFAy51::NifU::twin-Strep from *N. benthamiana*

*N. benthamiana* leaves infiltrated with SN166 and P19 were harvested 4 days post infiltration. Three (3.0) g leaf tissue was ground in 30 mL of 100 mM Tris-HCl pH 8.0, 150 mM NaCl, 5% (v/v) glycerol, 2 mM TCEP, 1% (w/v) PVP (average MW 40 kDa) and 0.1% Tween 20 using a mortar and pestle. The extract was centrifuged at 40,000 x *g* for 30 min at 10°C. The supernatant was loaded on a StrepTactinXT (IBA Lifesciences) column with a column volume of 2 mL equilibrated in 100 mM Tris-HCl pH 8.0, 150 mM NaCl, 2 mM TCEP (wash buffer). After loading, the column was washed with 20 mL wash buffer and the protein was eluted with 5 mL 100 mM Tris-HCl pH 8.0, 150 mM NaCl, 2 mM TCEP and 50 mM biotin. The eluate was concentrated using an Amicon® Ultra centrifugal filter (10 kDa MWCO). Samples from the supernatant, flow through and eluate were subjected to SDS-PAGE on a 4-12% NuPage SDS gel. Proteins were transferred to PVDF membranes with the iBlot dry blotting system (Thermo Fisher Scientific), washed with TBST and developed using the Strep-Tactin-HRp conjugate (IBA Lifesciences). The SDS gel was stained after blotting with SimplyBlue™ SafeStain (Thermo Fisher Scientific).

### 4.5 Construction of modified MIT v2.1 plasmids for function testing in *Escherichia coli*

First, MIT v2.1 was split into two parts for easier modification of the *nif* genes by PCR. The first half containing *nifHDKYENJ* was amplified with SbfI sites on either end (with oligos MIT_v2.1_SbfInifH_FW2 5’-*AACCTGCAGGTGACGTCTAAGAAAAGGAATATTCAGCAAT*-3’, and MIT_v2.1_SbfInifJ_RV2 5’-*AACCTGCAGGGCTAACTAACTAACCACGGACAAAAAACC*-3’) and ligated into pCR Blunt II TOPO (Thermo Fisher Scientific). The second half containing *nifBQFUSVWZM* was amplified with SbfI sites on either end (with oligos MIT_v2.1_SbfInifB_FW 5’-*AACCTGCAGGTACTCTAACCCCATCGGCCGTCTTA*-3’, and MIT_v2.1_SbfIori_RV 5’-*AACCTGCAGGTACGTAGCAATCAACTCACTGGCTC*-3’), digested with SbfI, and ligated back together. This religated plasmid, herein termed pB-ori, was used as a negative control for nitrogenase function testing. The positive control was constructed by ligating SbfI digested pCR Blunt II TOPO containing *nifHDKYENJ* and pB-ori. The scar9 extension (*ATGTCAACTCAAGTGGTGCGTAACCGC* coding for MSTQVVRNR) was added to the front of fw primers that bind to the start of the coding sequence for each *nif* gene, and rv primers were designed adjacent to the 5’ end of each *nif* gene that the scar9 was being added (primers listed in Suppl. Table 4). The amplified PCR product containing the scar9 extension in front of a given *nif* gene was ligated using ligation cycling reaction (LCR; de Kok et al., 2014). The other half of MIT v2.1 that was not modified was religated with the half with the scar9 extension via SbfI restriction sites. The plasmid ID and descriptions are listed in Suppl. Table 3.

### 4.6 Acetylene reduction assay

Acetylene reduction assays on *E. coli* transformed with control plasmids or modified MIT v2.1, along with controller plasmid N249 (Temme et al., 2012) were carried out according to Dilworth (1966) with the following modifications: Transformed JM109 cells were grown aerobically overnight at 37°C in LB medium with antibiotics to OD_600_ = 1.0. The cultures were resuspended in induction medium (25 g/L Na_2_HPO_4_, 3 g/L KH_2_PO_4_, 0.25 g/L MgSO_4_.7H_2_O, 1 g/L NaCl, 0.1 g/L CaCl_2._2H_2_O, 2.9 mg/L FeCl_3_, 0.25 mg/L Na_2_MoO_4._2H_2_O, 20 g/L sucrose, 0.015% serine, 0.5% casamino acids, 5 mg/L biotin, 10 mg/L para-aminobenzoic acid, 1 mM isopropyl β-D-thiogalactopyranoside (IPTG), transferred to air-tight culture flasks, and headspace sparged with argon gas. After 5 h incubation at 30°C, 200 rpm, pure C_2_H_2_ was injected into the headspace at 10% (vol.) and incubated for a further 18 h. Production of ethylene was measured by gas chromatography with flame ionisation detection (GC-FID) using a RT-Alumina Bond/MAPD column (30 m x 0.32 mm ID x 5 μm film thickness) with a 5 m particle trap column fitted to the detector end of an Agilent 6890N GC. Parameters for the GC-FID were as follows: the inlet and FID were set to 200°C, carrier gas (He) velocity at 35 cm/s, and isothermal oven temperature set to 120°C.

### 4.7 *E. coli* total protein extraction

Proteins were extracted from IPTG-induced *E. coli* JM109 containing modified MIT v2.1 plasmid as described above for the acetylene reduction assay using 8 M urea and 2% SDS in 100 mM Tris-HCl pH 8.5. Protein extracts were stored at -80°C prior to processing. Protein estimations were performed using the Bio-Rad microtiter Bradford protein assay (California, USA) according to the instructions provided (Bio-Rad version: Lit 33 Rev C) and measurements were made at 595 nm using a SpectraMax Plus. Bovine serum albumin (BSA) standard was used in the linear range 0.05 mg/mL to approximately 0.5 mg/mL. The BSA concentration was determined by high sensitivity amino acid analysis at Australian Proteomics Analysis Facility (Sydney, Australia).

### 4.8 *E. coli* tryptic digestion

Protein was subjected to filter-aided sample preparation (Wiśniewski et al., 2011). In brief, 100 µL (∼200 µg) of protein was diluted in 100 µL of 8 M urea, 100 mM Tris-HCl, pH 8.5 (UA buffer) and loaded onto a 10 kDa molecular weight cut-off (MWCO) centrifugal filter (Merck Millipore, Australia) and centrifuged at 20,800 x *g* for 15 min at 22°C. The filter (and protein >10 kDa) was washed with 200 µL of UA buffer and centrifuged at 20,800 x *g* for 15 min at 22°C. To reduce the protein on the filter, dithiothreitol (50 mM, 200 µL) was added and the solution incubated at room temperature for 50 min with shaking. The filter was washed with two 200 µL volumes of UA buffer with centrifugation (20,800 x *g*, 15 min). For cysteine alkylation, iodoacetamide (IAM) (100 µL, 50 mM IAM in UA buffer) was added and incubated in the dark for 30 min at 22°C before centrifugation (20,800 x *g*, 15 min) and washed with two 200 µL volumes of UA buffer with centrifugation (20,800 x *g*, 15 min) followed by two subsequent wash/centrifugation steps with 200 µL of 50 mM ammonium bicarbonate. The trypsin (sequencing grade, Promega, Alexandria, Australia) solution (200 µL, 20 µg/mL (4 µg) in 50 mM ammonium bicarbonate and 1 mM CaCl_2_) was loaded onto the filter and incubated for 18 h at 37°C in a wet chamber. The tryptic peptides were collected by centrifugation (20,800 x *g*, 15 min) followed by an additional wash with 200 µL of 50 mM ammonium bicarbonate. The combined filtrates were lyophilized and stored at -20°C.

### 4.9 Global proteomic profiling

The digested peptides were reconstituted in 50 µL of 1% formic acid (FA) and chromatographic separation (4 μL) on an Ekspert nanoLC415 (Eksigent, Dublin, CA, U.S.A.) directly coupled to a TripleTOF 6600 liquid chromatography tandem mass spectrometry (LC-MS/MS, SCIEX, Redwood City, CA, USA). The peptides were desalted for 5 min on a ChromXP C18 (3 μm, 120 Å, 10 mm × 0.3 mm) trap column at a flow rate of 10 μL/min 0.1% FA, and separated on a ChromXP C18 (3 μm, 120 Å, 150 mm × 0.3 mm) column at a flow rate of 5 μL/min at 30°C. A linear gradient from 3-25% solvent B over 68 min was employed followed by: 5 min from 25% B to 35% B; 2 min 35% B to 80% B; 3 min at 80% B, 80-3% B, 1 min; and 8 min re-equilibration. The solvents were: (A) 5% dimethylsulfoxide (DMSO), 0.1% formic acid (FA), 94.9% water; (B) 5% DMSO, 0.1% FA, 90% acetonitrile, 4.9% water. The instrument parameters were: ion spray voltage 5500 V, curtain gas 25 psi, GS1 15 psi and GS2 15 psi, heated interface 150°C. Data were acquired in information-dependent acquisition mode comprising a time-of-flight (TOF)-MS survey scan followed by 30 MS/MS, each with a 40 ms accumulation time. First stage MS analysis was performed in positive ion mode, mass range *m/z* 400-1250 and 0.25 s accumulation time. Tandem mass spectra were acquired on precursor ions >150 counts/s with charge state 2-5 and dynamic exclusion for 15 s with a 100 ppm mass tolerance. Spectra were acquired over the mass range of *m/z* 100-1500 using the manufacturer’s rolling collision energy based on the size and charge of the precursor ion. Protein identification was undertaken using ProteinPilot™ 5.0 software (SCIEX) with searches conducted against the *E. coli* subset of the Uniprot-SwissProt database (2018/08) appended with a custom nitrogenase (Nif+Mit2Nif) database including the control chloramphenicol acetyltransferase (CAT/P62577) and a contaminant database (Common Repository of Adventitious Proteins). The total number of proteins in the custom database was 5410.

### 4.10 Identification of prototypic peptides for nitrogenase proteins in *E. coli*

From the identified peptides, two NifM peptides (DAFAPLAQR and DYLWQQSQQR) that were fully tryptic, contained no unusual cleavages and/or modifications and showed high response in the MS (as judged by peak intensity) were selected for multiple reaction monitoring scanning to confirm the detection of the nitrogenase (NifM) proteins in the *E. coli* JM109 expressions.

### 4.11 Targeted liquid chromatography – multiple reaction monitoring – mass spectrometry (LC-MRM-MS)

Reduced and alkylated tryptic peptides (5 μL) were chromatographically separated on a Kinetex C18 column (2.1 mm x 100 mm, Phenomenex) using a linear gradient of 5–45% acetonitrile (in 0.1% formic acid) over 10 min at a flow rate of 400 μL/min. The eluent from the Shimadzu Nexera UHPLC was directed to a QTRAP 6500 mass spectrometer (SCIEX) equipped with a TurboV ionisation source operated in positive ion mode for data acquisition and analysis. The MS parameters were as follows: ion spray voltage, 5500 V; curtain gas, 35; GS1, 35; GS2, 40; source temperature, 500°C; declustering potential, 70 V; and entrance potential, 10 V. Peptides were fragmented in the collision cell with nitrogen gas using rolling collision energy dependent on the size and charge on the size and charge of the precursor ion. Relative quantitation using scheduled multiple reaction monitoring (MRM) scanning experiments (MRM transition peptide information provided in Supplementary Table 1) with a 40 second detection window around the expected retention time (RT) and a 0.3 second cycle time. Data were acquired using Analyst v1.7 software. Peak areas of four MRM transitions were integrated using Skyline (MacLean, Bioinformatics 2010) wherein all transitions were required to co-elute with a signal-to-noise (S/N) > 3 and intensity >1000 counts per second (cps) for detection.

### 4.12 Identification of peptides for chloramphenicol acetyltransferase protein

Chloramphenicol acetyltransferase (CAT/P62577) enzyme is an effector of chloramphenicol resistance in bacteria and is expressed in all *E. coli* JM109 transformed with unmodified or modified MIT v2.1. This protein was selected as a control for protein expression. Three peptides (four transitions/peptide) were selected to screen the expression of CAT (ITGYTTVDISQWHR, LMNAHPEFR, and YYTQGDK).

## 6 Author Contributions

SO, CG, RA, CW conceived the project and designed the experiments. SO, CG, RA, AM, DH, VG, EJ, KB, MC conducted the experiments. All authors contributed to writing the manuscript.

## 8 Acknowledgments

We thank Rob Defeyter, Trevor Rapson and Xue-Rong Zhou for their critical reviews of this article.

## 12 Supplementary material

**Suppl. Fig. 1:**
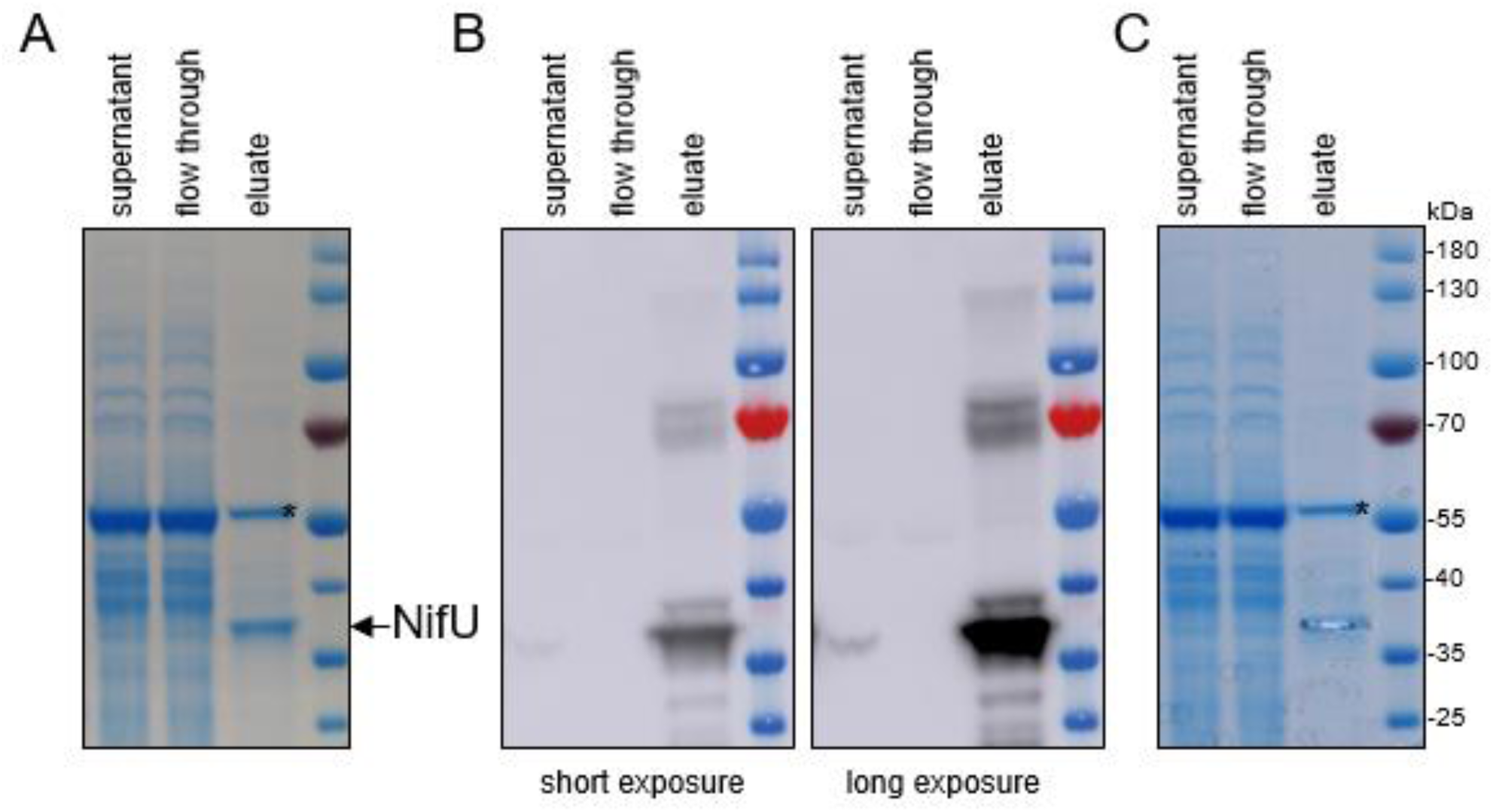
Purification of NifU from plant leaves and sample preparation for proteomic analysis. **(A)** Coomassie stain of the supernatant, flow through and eluate from the StrepTactin purification. A contaminating band (*) was observed, most likely corresponding to Rubisco (large chain). **(B)** Western blot analysis of the same samples with a StrepTactin-HRP conjugate antibody. **(C)** Coomassie gel after excision of the NifU gel slice for proteomic analysis.

**Suppl. Fig. 2:**
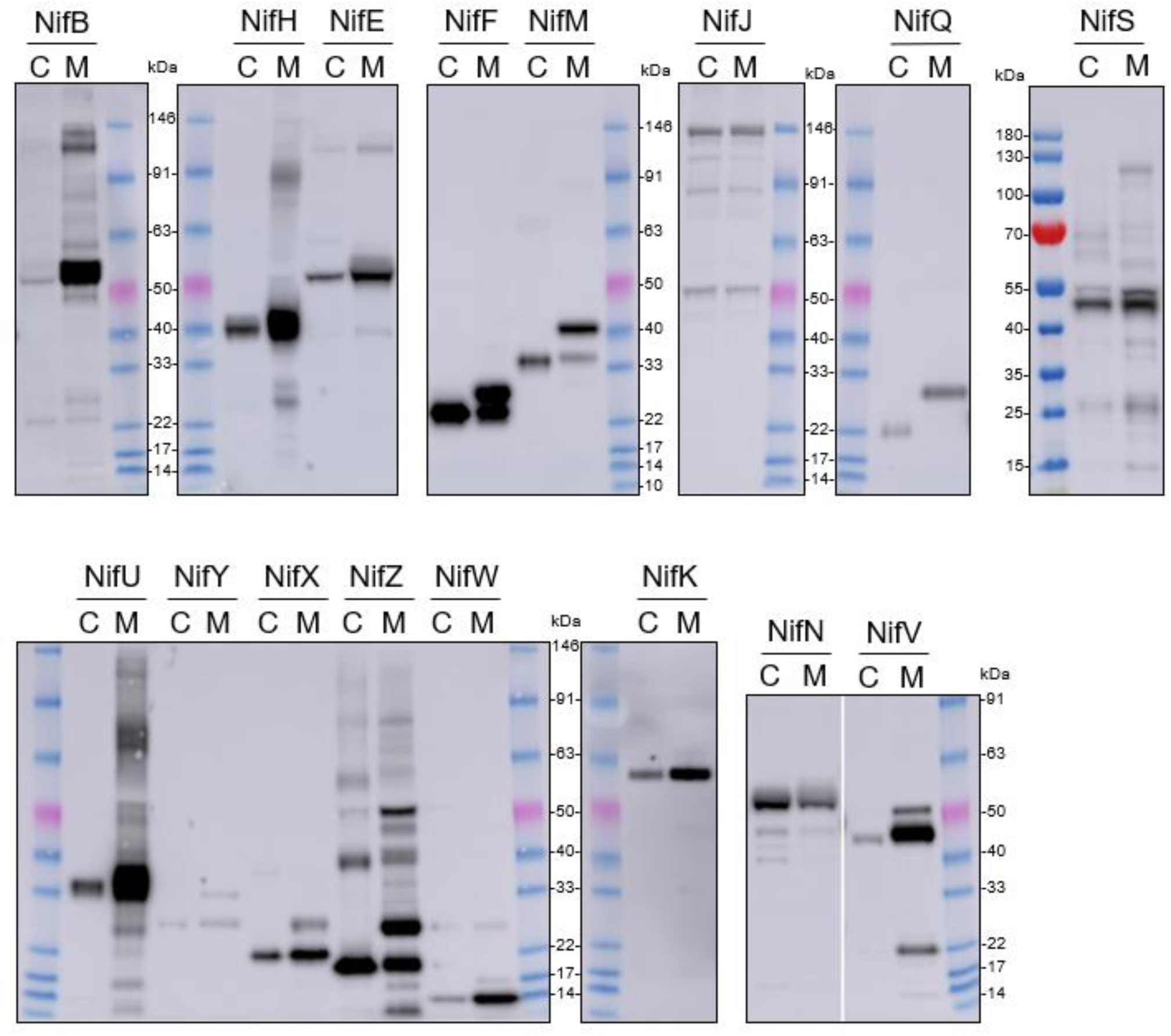
Assessment of MTP cleavage of pFAγ51::Nif::HA proteins. Whole blot images of the Western blot analysis of individual pFAγ51::Nif::HA, pFAγ51::HA::NifK, 6×His::Nif::HA and 6×His::HA::NifK proteins transiently expressed in *Nicotiana benthamiana* leaf. C, cytoplasmic expression; M, mitochondrially targeted. Due to considerable variation in abundance of mitochondrially located proteins and cytoplasmic equivalents, mitochondrially located NifB, NifE, NifF, NifH, NifK, unprocessed NifM, NifN, NifU, NifV, NifW, NifX and NifZ in these images are overexposed. Cytoplasmic NifF and NifZ are also overexposed.

**Suppl. Fig. 3:**
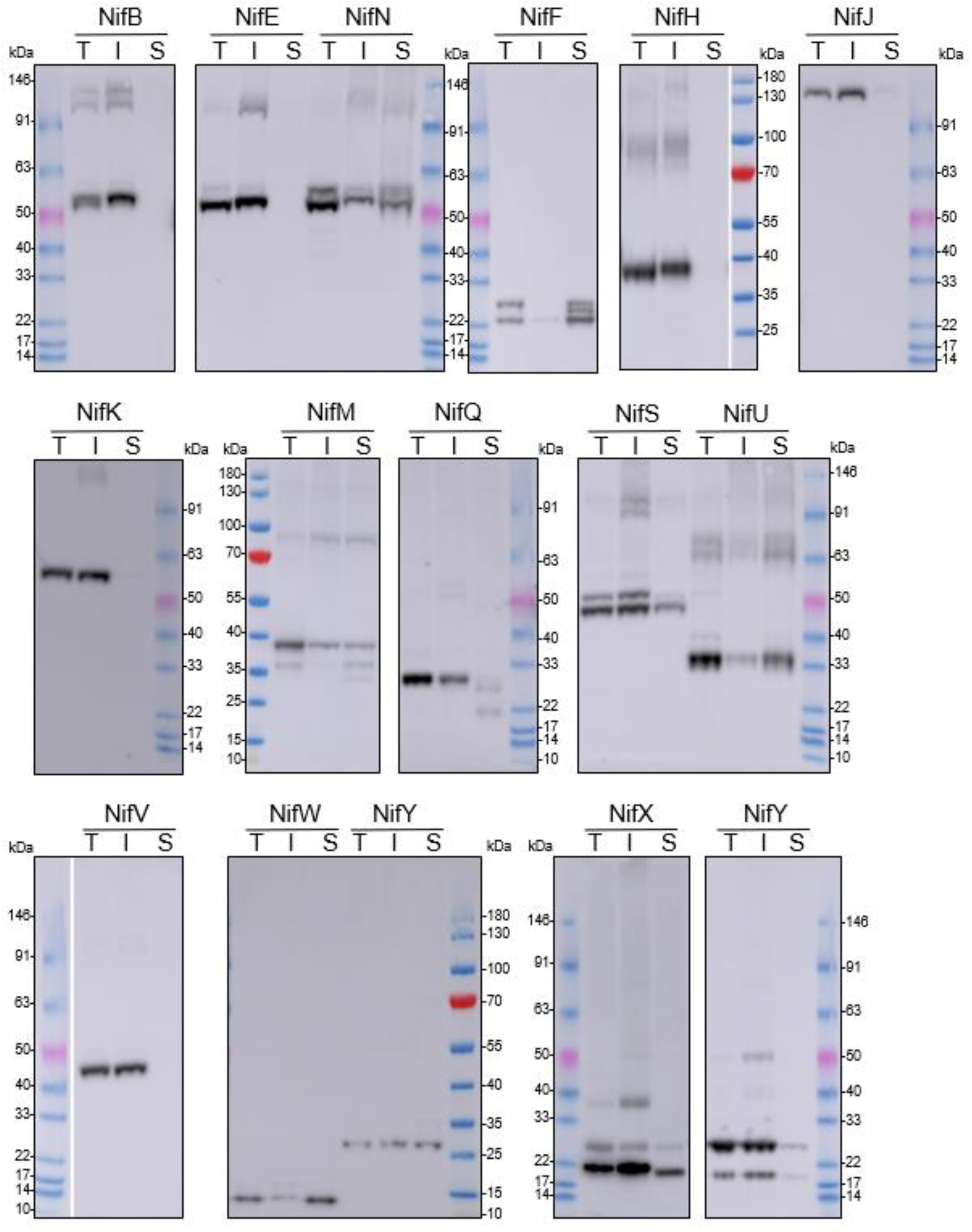
Whole blot images of the Western blot analysis (α-HA) of individual pFAγ51::Nif::HA and pFAγ51::HA::NifK proteins transiently expressed in *N. benthamiana* leaf. T, total protein; I, insoluble fraction; S, soluble fraction.

**Suppl. Fig. 4:**
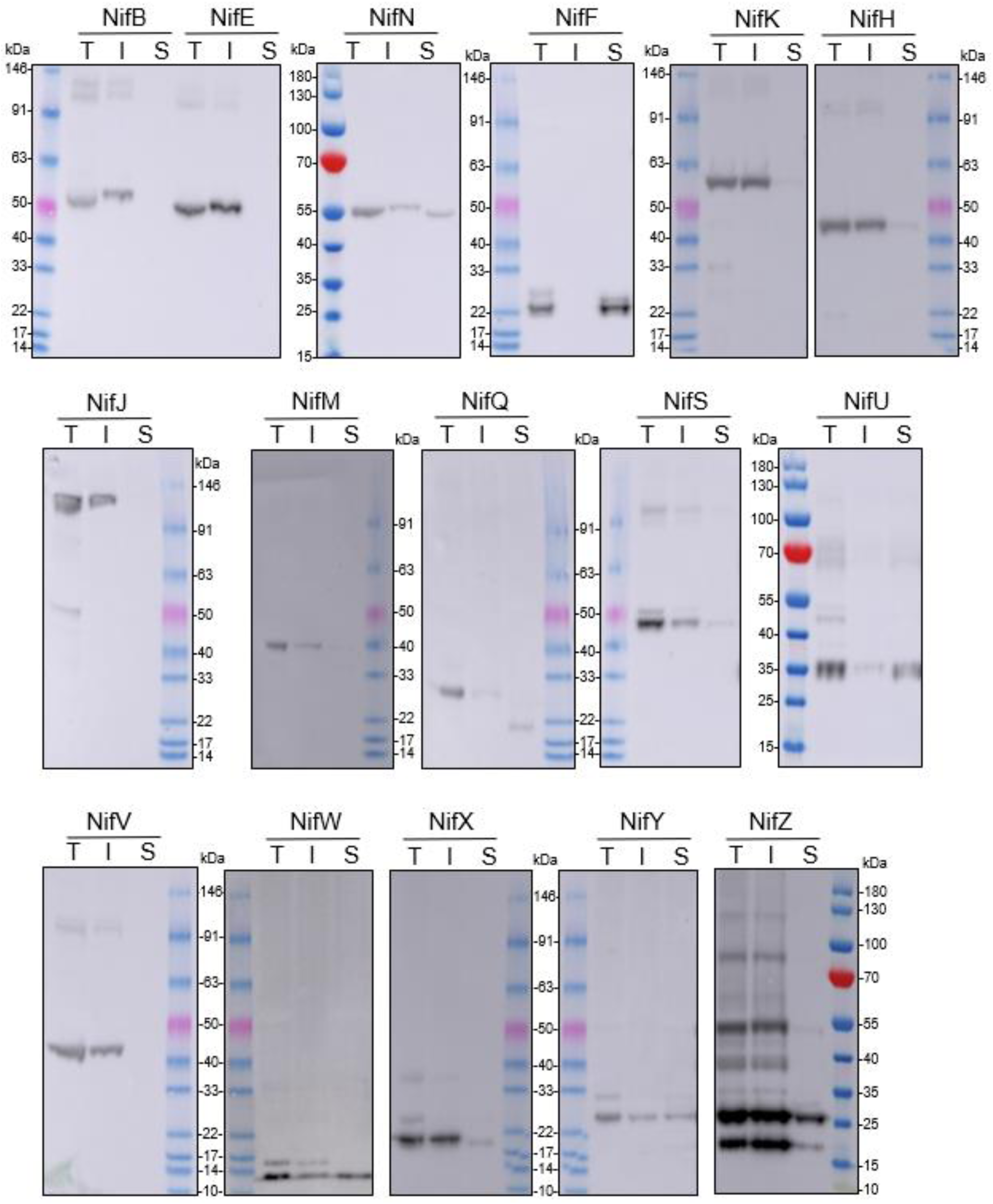
Whole blot images of Western blot analysis (α-HA) of individual pFAγ51::Nif::HA and pFAγ51::HA::NifK proteins transiently expressed in *N. benthamiana* leaf. Proteins were extracted under anaerobic conditions. T, total protein; I, insoluble fraction; S, soluble fraction.

**Suppl. Fig. 5:**
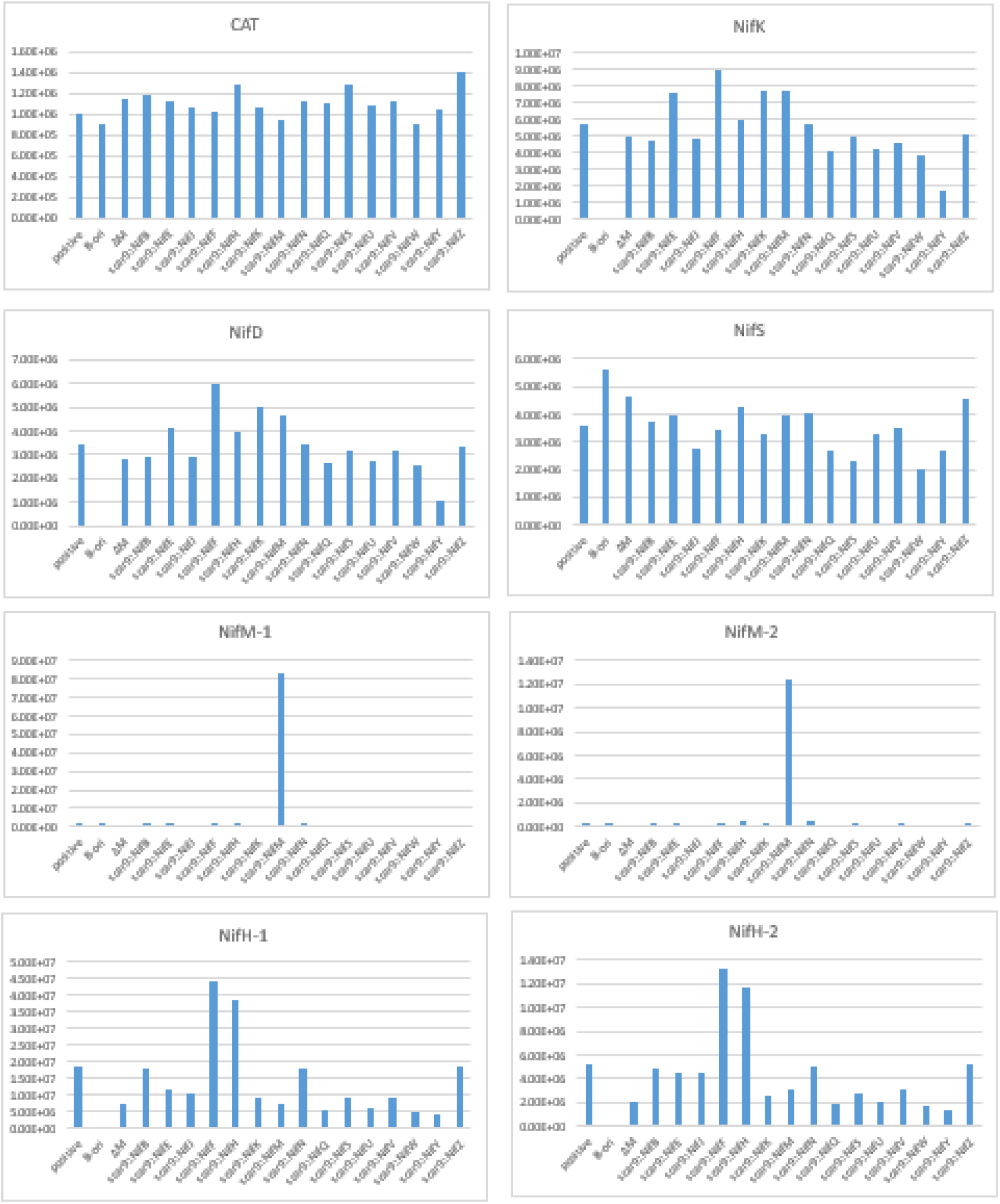
Relative expression levels of chloramphenicol acetyltransferase peptide YYTQGDK (CAT) and Nif peptides in *E. coli* JM109 expressing unmodified and modified MIT v2.1 plasmids. Positive, positive control - unmodified MIT v2.1; pB-ori, negative control containing *nifBQFUSVWZM*; ΔM, MIT V2.1 with nifM deleted. Peptide information for CAT and Nif proteins are provided in Supplementary Table 1.

**Suppl. Table 1:** Multiple reaction monitoring transitions of peptides for targeted liquid chromatography – multiple reaction monitoring – mass spectrometry.

| Protein | Peptide <sup>a</sup> | RT<br>(min) <sup>b</sup> | Q1<br>*m/z* <sup>b</sup> | *z* <sup>b</sup> | Q3<br>*m/z* <sup>a</sup> | Fragment | CE <sup>c</sup> |
| --- | --- | --- | --- | --- | --- | --- | --- |
| CAT | YYTQGDK | 1.0 | 437.70 | 2+ | 319.16 | y3+ | 20.45 |
|  |  |  |  |  | 447.22 | y4+ | 20.45 |
|  |  |  |  |  | 548.26 | y5+ | 20.45 |
|  |  |  |  |  | 711.33 | y6+ | 20.45 |
| NifK | EALTVDPAK | 3.5 | 472.25 | 2+ | 315.20 | y3+ | 22.14 |
|  |  |  |  |  | 430.22 | y4+ | 22.14 |
|  |  |  |  |  | 529.29 | y5+ | 22.14 |
|  |  |  |  |  | 630.34 | y6+ | 22.14 |
| NifD | SMNYIAR | 3.2 | 427.71 | 2+ | 246.15 | y2+ | 19.96 |
|  |  |  |  |  | 359.24 | y3+ | 19.96 |
|  |  |  |  |  | 522.30 | y4+ | 19.96 |
|  |  |  |  |  | 636.34 | y5+ | 19.96 |
| NifS | QVYLDNNATTR | 3.2 | 647.82 | 2+ | 676.33 | y6+ | 30.74 |
|  |  |  |  |  | 791.36 | y7+ | 30.74 |
|  |  |  |  |  | 904.44 | y8+ | 30.74 |
|  |  |  |  |  | 1067.51 | y9+ | 30.74 |
| NifH-1 | VMIVGC(cam)DPK | 3.9 | 509.75 | 2+ | 576.24 | y5+ | 23.97 |
|  |  |  |  |  | 675.31 | y6+ | 23.97 |
|  |  |  |  |  | 788.39 | y7+ | 23.97 |
|  |  |  |  |  | 231.11 | b2+ | 23.97 |
| NifH-2 | LGGLIC(cam)NSR | 3.8 | 495.26 | 2+ | 536.22 | y4+ | 23.26 |
|  |  |  |  |  | 649.30 | y5+ | 23.26 |
|  |  |  |  |  | 876.43 | y8+ | 23.26 |
|  |  |  |  |  | 228.13 | b3+ | 23.26 |
| NifM-1 | DAFAPLAQR | 4.68 | 494.76 | 2+ | 584.35 | y5+ | 23.24 |
|  |  |  |  |  | 655.39 | y6+ | 23.24 |
|  |  |  |  |  | 802.46 | y7+ | 23.24 |
|  |  |  |  |  | 334.14 | b3+ | 23.24 |
| NifM-2 | DYLWQQSQQR | 4.47 | 676.32 | 2+ | 646.33 | y5+ | 32.14 |
|  |  |  |  |  | 774.39 | y6+ | 32.14 |
|  |  |  |  |  | 960.46 | y7+ | 32.14 |
|  |  |  |  |  | 279.10 | b2+ | 32.14 |

a. The peptide sequence is represented by single amino acid code. C (cam) refers to carbamidomethylation of cysteine.
b. RT, retention time (min); Q1 $m/z$ , precursor ion mass-to-charge ratio; z, charge state; Q3 $m/z$ , fragment ion $m/z$ ; CE, collision energy in V.
c. Collision energy settings were calculated ( $CE = \text{slope } (0.049) \times (\text{precursor } m/z) + \text{intercept}$ $(-1.0)$ ) for a 6500 QTRAP mass spectrometer (AB SCIEX, Redwood City, USA).

**Suppl. Table 2:** Designations and descriptions of plasmids constructed to test mitochondrial targeting efficiency and protein solubility of Nif proteins transiently expressed in *Nicotiana benthamiana*. The protein size (kDa) was calculated using the average molecular weight of an amino acid as 0.11 kDa.

| Plasmid ID | Description | calculated protein size<br>(kDa) |
| --- | --- | --- |
| SN192 | pFAγ51::NifB::HA | 59 |
| SN38 | pFAγ51::NifE::HA | 57 |
| SN138 | pFAγ51::NifF::HA | 26 |
| SN27 | pFAγ51::NifH::HA | 39 |
| SN139 | pFAγ51::NifJ::HA | 135 |
| SN140 | pFAγ51::HA::NifK | 64 |
| SN30 | pFAγ51::NifM::HA | 36 |
| SN39 | pFAγ51::NifN::HA | 58 |
| SN141 | pFAγ51::NifQ::HA | 25 |
| SN31 | pFAγ51::NifS::HA | 51 |
| SN32 | pFAγ51::NifU::HA | 37 |
| SN142 | pFAγ51::NifV::HA | 49 |
| SN143 | pFAγ51::NifW::HA | 16 |
| SN144 | pFAγ51::NifX::HA | 24 |
| SN145 | pFAγ51::NifY::HA | 31 |
| SN146 | pFAγ51::NifZ::HA | 23 |
| SN201 | 6×His::NifB::HA | 54 |
| SN203 | 6×His::NifE::HA | 52 |
| SN204 | 6×His::NifF::HA | 21 |
| SN205 | 6×His::NifH::HA | 34 |
| SN206 | 6×His::NifJ::HA | 130 |
| SN72 | 6×His::HA::NifK | 59 |
| SN207 | 6×His::NifM::HA | 31 |
| SN208 | 6×His::NifN::HA | 53 |
| SN209 | 6×His::NifQ::HA | 20 |
| SN210 | 6×His::NifS::HA | 46 |
| SN211 | 6×His::NifU::HA | 32 |
| SN212 | 6×His::NifV::HA | 44 |
| SN213 | 6×His::NifW::HA | 11 |
| SN214 | 6×His::NifX::HA | 19 |
| SN215 | 6×His::NifY::HA | 26 |
| SN216 | 6×His::NifZ::HA | 18 |
| SN149 | pFAγ77::NifM::HA | 39 |
| SN151 | pFAγ77::NifQ::HA | 28 |
| SN247 | pFAγ51m::NifS::HA | 51 |
| SN166 | pFAγ51::NifU::twin-Strep | 39 |

**Suppl. Table 3:** Designations and descriptions of plasmids constructed for bacterial function testing in *Escherichia coli*.

| Plasmid ID | Description |
| --- | --- |
| pSO003 | MIT v2.1 |
| pSO006 | scar9::NifB |
| pSO009 | scar9::NifD |
| pSO026 | scar9::NifE |
| pSO032 | scar9::NifF |
| pSO012 | scar9::NifH |
| pSO028 | scar9::NifJ |
| pSO029 | scar9::NifK |
| pSO038 | scar9::NifM |
| pSO027 | scar9::NifN |
| pSO031 | scar9::NifQ |
| pSO034 | scar9::NifS |
| pSO033 | scar9::NifU |
| pSO035 | scar9::NifV |
| pSO036 | scar9::NifW |
| pSO030 | scar9::NifY |
| pSO037 | scar9::NifZ |
| pSO051 | $\Delta nifM$ |
| pSO013 | NifK::HA |
| pSO004 | pB-ori<br>($\Delta NifHDKENJ$) |

**Suppl. Table 4:**
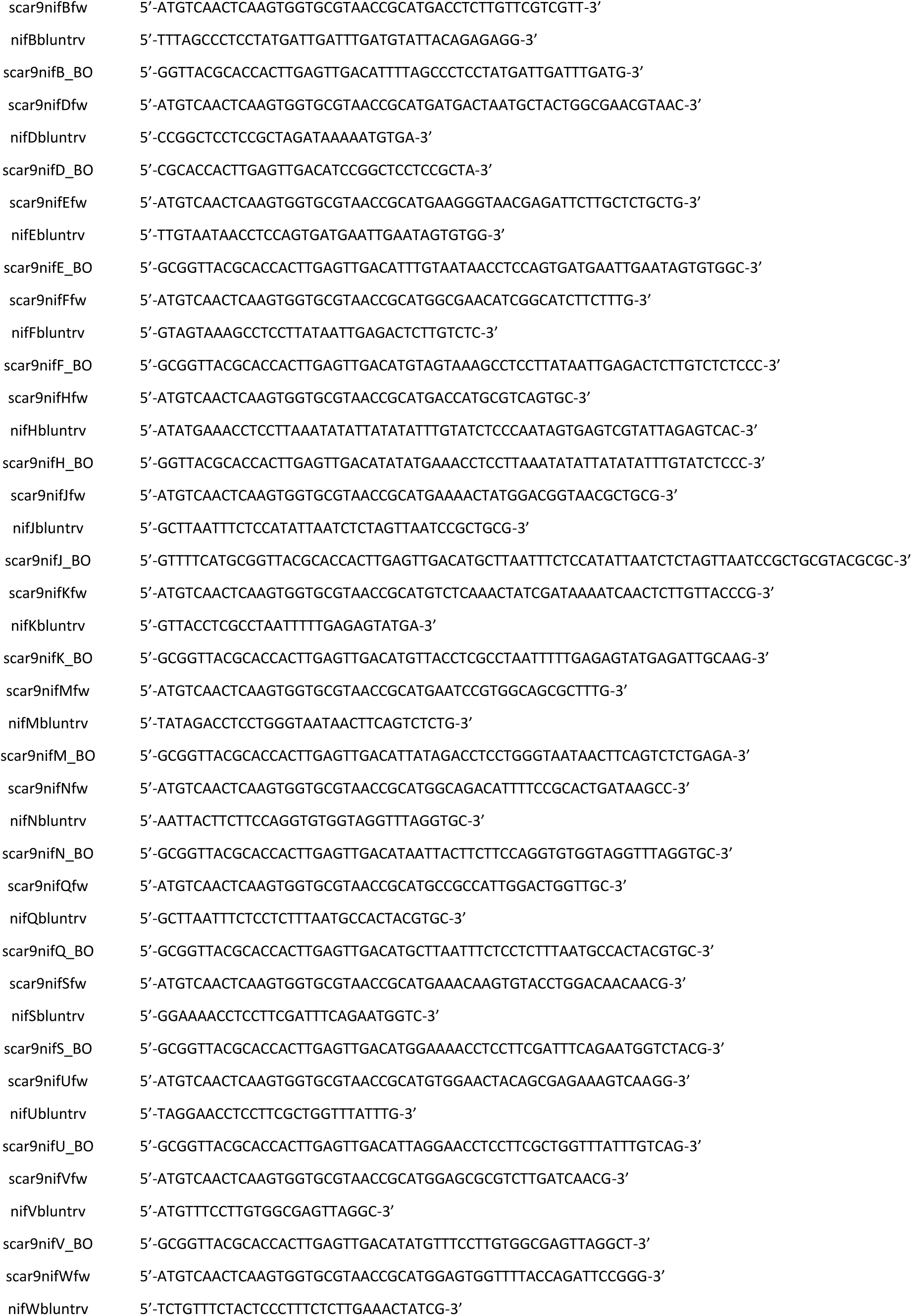
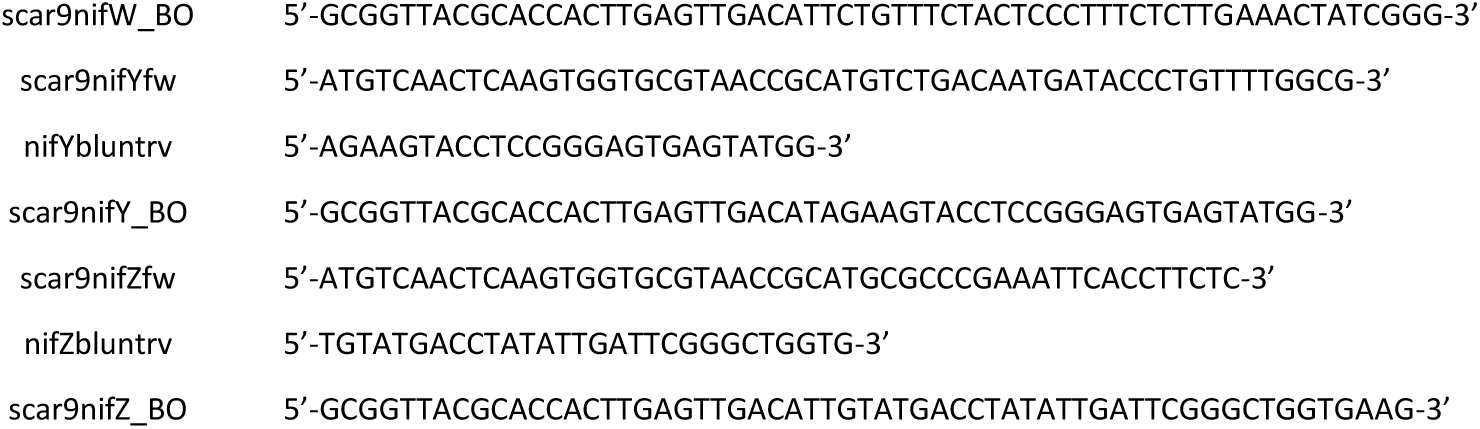
Primers used to add the scar9 peptide onto the N-terminus of each Nif protein via translational fusion. BO, bridging oligo.

